# How Bias Shapes the Leaderboard: Scoring Function Performance Under Scrutiny

**DOI:** 10.64898/2026.07.27.740774

**Authors:** David Graber, Jakub Kopko, Peter Stockinger, Ravidu Nakandalage, Bernd Kuhn, Siddhartha Mishra

## Abstract

Structure-based scoring functions leveraging machine learning have recently demonstrated superior performance over classical scoring functions, particularly on virtual screening benchmarks. However, due to the fundamental differences between their underlying model principles and architectures, it remains unclear to what extent performance stems from an understanding of molecular binding or from exploitation of systemic biases. Thus, disentangling the factors underlying benchmark performance is essential for determining whether a scoring function will generalize to novel chemical space and succeed in prospective drug discovery.

To address this need, we present a case study investigating the nature and impact of systemic biases on benchmark comparisons between different scoring function paradigms. By systematically analyzing the evaluation workflows of prominent models, we reveal ‘pocket bias’, a form of spatial coordinate frame leakage arising from static binding pocket extraction, which artificially inflates benchmark performance. To progressively eliminate these sources of bias, we benchmarked two selected graph neural network scoring functions against two minmalist machine learning models and a classical scoring function under four increasingly stringent evaluation levels, successively removing pocket bias, reducing structural data leakage, and finally evaluating on out-of-distribution (OOD) protein targets.

Upon removal of pocket bias and structural data leakage, the performance of all machine learning models dropped substantially. When evaluated on out-of-distribution protein families, the classical baseline AutoDock Vina outperformed the machine learning models in five of seven virtual screening tasks and dominated the docking power evaluation.

Our findings indicate that benchmark performance can be heavily shaped by evaluation design and dataset artifacts, potentially overshadowing algorithmic improvements. While the tested machine learning models remain heavily dependent on encountering familiar data distributions to achieve competitive results, AutoDock Vina demonstrated superior generalization capacity on OOD targets. This work underscores the critical need for rigorous, artifact-free benchmarking protocols to guide the development of truly prospective machine learning models for virtual screening.

---

Structure-based virtual screening (VS) is an important and established pillar of modern drug discovery [1]. It represents the strategic application of computational filters leveraging 3D protein structures to prioritize chemical compounds from large libraries before committing to expensive and time-consuming in vitro assays [1, 2]. The primary objective of VS is the identification of small molecules that exhibit high binding affinity to a specific protein target. Typically, the process begins with library preparation, where ligands are filtered for desired molecular properties and processed into three-dimensional conformations of the relevant protomers. This involves conformational sampling, the algorithmic exploration of a ligand’s degrees of freedom within the binding site, to identify plausible interaction poses. In the subsequent scoring step, a scoring function evaluates the binding affinity of each generated conformation [1, 3].

The limited accuracy of scoring functions (SFs) has historically been the bottleneck of structure-based VS pipelines [3, 4]. For many years, the field relied on classical SFs, which are generally categorized as physics-based [5, 6], empirical [7–9], or knowledge-based [10, 11]. Recently, there was a growing interest in Machine Learning Scoring Functions (ML-SFs) and their potential to complement or replace traditional approaches. In this paradigm, 3D conformations and target-ligand interactions are evaluated by a Machine Learning (ML) model that has learned complex structural relationships directly from large structural datasets [4, 12, 13]. These ML-SFs, ranging from Random Forests (RF) [14] to deep learning architectures like Graph Neural Networks (GNNs) [15–19], have consistently been reported to possess superior predictive accuracy compared to classical SFs [12, 15, 20].

Effective scoring of 3D binding poses necessitates a rigorous framework for evaluation [21]. In this context, the field commonly differentiates between the specific “powers” of a SF to solve distinct aspects of the VS task: docking, screening, and scoring powers [22]. An ideal SF exhibits high performance across all three tasks simultaneously.

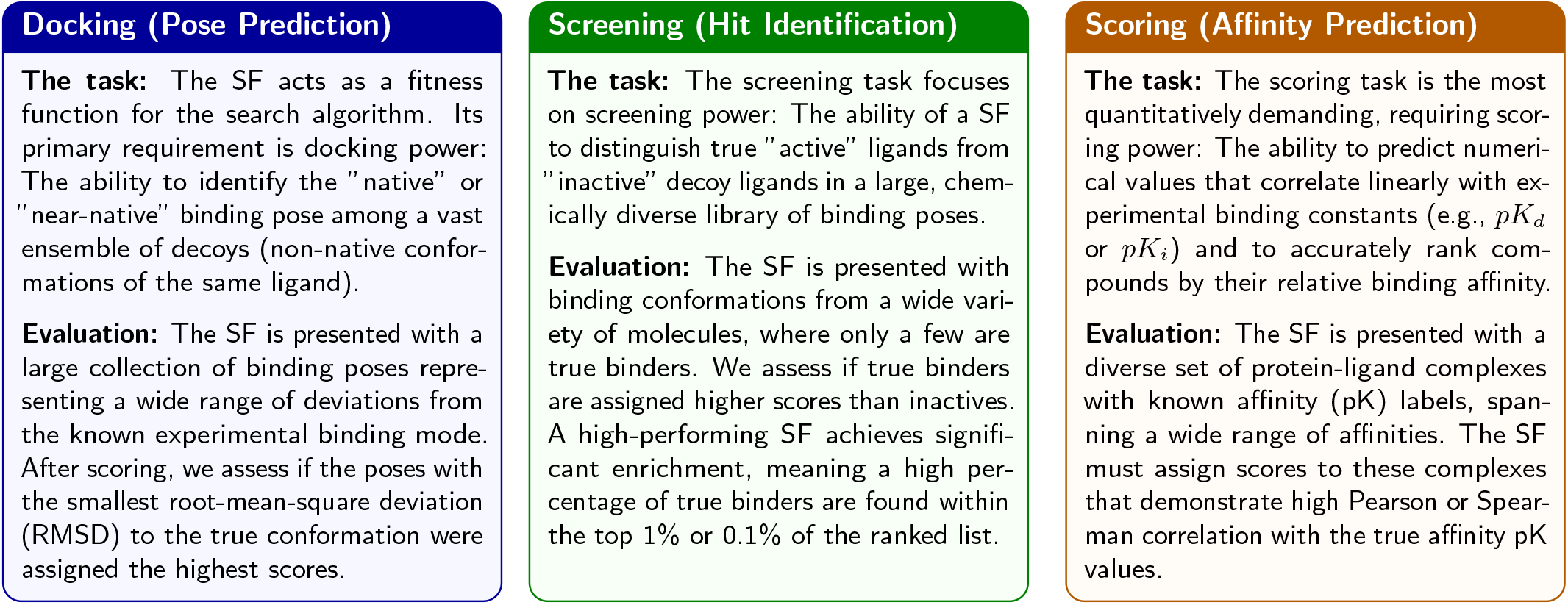

Considering the difficulty of these tasks, it is remarkable that high performance has been reported by several recent ML-SFs across docking, screening, and scoring tasks of the CASF2016 benchmark [22], outperforming all classical SFs [16–18].

The primary difficulty in interpreting these results lies in the fundamental differences between the compared functions. While classical SFs rely on explicit physical parameters to calculate binding energies [1], data-driven ML and deep learning models operate as black boxes. Consequently, it often goes undetected when a ML model achieves accurate benchmark predictions due to noncausal bias, a phenomenon where models exploit correlative patterns in the data that do not represent any biologically relevant binding mechanism. A classic manifestation of this bias is seen in ligand-only models, which have been shown to perform remarkably well across benchmarks despite completely ignoring the protein target structure [23–25].

Furthermore, ML models can build complex internal representations that are able to “memorize” training samples [26–29]. This tendency to memorize has been demonstrated repeatedly, particularly regarding models trained on the PDBbind database [30, 31] and evaluated on the CASF2016 benchmark, where a train-test data leakage seems to exist [15, 23, 24, 32–37]. Consequently, CASF2016 metrics often overestimate a model’s true generalization capability due to structural similarities between the training and test sets [15, 23, 33]. Previous studies have shown that model performance degrades when this similarity is reduced during OOD evaluation [19, 24, 38, 39]. To address these problems, recent efforts have focused on compiling filtered training or test datasets, such as PDBbind CleanSplit [15], Peptide Holdout [24], 0 Ligand Bias [24] and LeakProof PDBbind [35], which aim to eliminate train-test data leakage [19].

In addition to these biases linked to distributional overlap, we found that systemic biases can be introduced involuntarily during data preparation. Specifically, during the encoding of docking poses into graphs for scoring with GNNs. Such encoding biases can provide the GNN model with unintended cues, resulting in an information “leakage” occurring during the conversion of 3D structures into graphs that facilitates accurate scoring of molecular interactions in the benchmark.

Therefore, we believe a key challenge in evaluating modern structure-based SFs is not merely measuring raw benchmark performance, but understanding the origin of that performance and verifying whether it is based on generalizable capabilities. While high metrics are frequently reported for ML-SFs, it often remains unclear to what extent these results reflect a genuine understanding of molecular binding or the exploitation of systemic biases. Without a systematic analysis of how factors like data proximity and encoding biases influence outcomes, the perceived superiority of one model over another remains fundamentally opaque. Identifying these drivers of accuracy is essential for determining whether a model will truly generalize to novel chemical space or fail in a prospective drug discovery setting.

The primary objective of this work is to illustrate the profound impact of these biases on the performance and comparative evaluation of structure-based SFs. Rather than attempting an exhaustive benchmarking of all available methods, we employ a selection of state-of-the-art ML and classical SFs as a case study to demonstrate how performance metrics on the CASF2016 benchmark fluctuate when specific biases are isolated.

In addition to the physics-based classical SF AutoDock Vina [8], we evaluate two recent GNN-based ML-SFs: GEMS [15] and GenScore [16]. These models were selected due to their strong performance on the CASF2016 subtasks and their shared training database, which enables a controlled comparison free from effects of varying training data. GEMS is a relatively simple GNN regressor trained for the scoring task, where it has demonstrated excellent CASF2016 benchmark performance, even under evaluation setups with minimized train-test data leakage. GenScore employs a Mixture Density Network (MDN) trained to predict pairwise distances between protein residues and ligand atoms. It was selected as it represents one of the first ML models to simultaneously outperform classical SFs across scoring, docking, and screening power.

In the following sections, we compare the performance of the aforementioned models across four progressive levels of evaluation, where systemic advantages often afforded to ML-SFs, such as the inclusion of target-proximal data or biased structural encoding, are progressively removed. Through this approach, we show how performance metrics fluctuate when these artifacts are eliminated and demonstrate that benchmark performance in ML-SFs is heavily influenced by experimental design choices rather than algorithmic superiority. Ultimately, we hope these insights help better contextualize obtained performance metrics and encourage the field to adopt more rigorous standards for the comparison of ML and classical SFs.

**Level 0: Original GNN Models (Reproduction)**

We evaluate the models using their original architectures and training protocols, establishing a baseline reproduction of their performance via 5-fold cross-validation.

**Level 1: Removal of Encoding Bias**

We eliminate an encoding bias in GenScore that inflates its docking and screening power metrics, establishing a fair performance comparison of the two GNN-based models trained on PDBbind.

**Level 2: Removal of Structural Data Leakage**

We remove a train-test overlap between PDBbind and CASF2016 and thus remove the advantage of closely related complexes in the training datasets of the ML-based models.

**Level 3: Evaluation Under Distribution Shift (OOD Datasets)**

We test the models on Out-of-Distribution (OOD) datasets featuring novel protein families, assessing their real-world generalizability and ability to extrapolate physical binding principles.

## Results

We retrained GEMS and GenScore using the publicly available code and data, strictly following the authors instructions. To gain insight into the emergence of the predictive accuracy of GEMS and GenScore, we tracked their scoring, docking, and screening performance at regular intervals throughout the training process, allowing us to visualize how these metrics manifest over successive training epochs.

To contextualize the results obtained with ML-SFs, we establish several baselines. As main baseline across the scoring, docking and screening tasks, we use the physics-based classical SF AutoDock Vina [8]. As an additional baselines for the scoring power task, we implement two sequence-based models based on the classical machine learning algorithms XGBoost and *K*-Nearest Neighbors (KNN) trained on language model (LM) embeddings extracted from MolFormer (for ligands) [40] and ESM2 (for proteins) [41]. The KNN baseline serves as an intentional instance-based baseline, designed to demonstrate the scoring performance achievable purely through training dataset memorization.

When comparing different ML models, it is essential to eliminate confounding variables introduced by differing evaluation protocols. To ensure a strictly fair comparison, we train and evaluate all ML-based models using an identical 5-fold cross-validation pipeline and report mean and standard deviation of the benchmark performance across the five resulting models. Because each training fold contains only 4/5 of the training data, which is a slight reduction compared to some original published implementation, the absolute performance metrics reported herein may be marginally lower than the published values. This deliberate standardization allows for statistically rigorous comparisons that reflect algorithmic superiority. A comprehensive justification of this methodological trade-off is detailed in the Methods section.

### Level 0: Original GNN Models (Reproduction)

In initial reproduction experiments, both GEMS and GenScore were retrained on the full PDBbind database and evaluated on the CASF2016 benchmark, as in the original studies. Five independent instances were trained for each architecture (5-fold CV), with each run continuing until the author-implemented early stopping mechanism was triggered. This resulted in short training runs (100–200 epochs) for GEMS and much longer training runs (1000-1400 epochs) for GenScore (Fig. 1).

**Figure 1:**
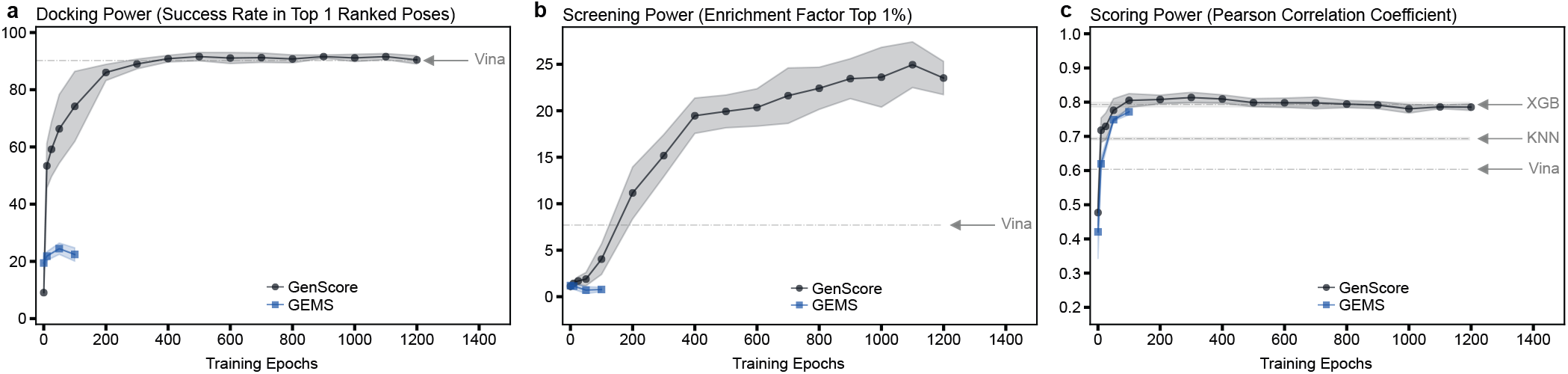
Evolution of CASF2016 metrics of GEMS and GenScore during model training: Comparison of **a)** docking, **b)** screening and **c)** scoring performance of GenScore, GEMS and AutoDock Vina. The scoring power plot includes two sequence-based classical machine learning models (XGBoost and *K*-Nearest-Neighbor) trained on language model embeddings of the protein-ligand complexes. In all plots, the x-axis shows training progress (epochs), while the y-axis displays the Pearson Correlation coefficient (↑, for scoring), the success rate (↑, for docking) and the enrichment factors (↑, for screening). For machine learning-based models, each line represents the mean performance across five cross-validation (CV) folds, with shaded uncertainty regions (±1 standard deviation) indicating the variability in performance across the five training runs. To maintain statistical rigor when different model instances terminated at different epoch numbers, only epochs with at least three active CV models contributing performance data are plotted.

In the **docking power** and **screening power** tasks (Fig. 1a,b), GEMS, which was optimized specifically for the scoring task, shows poor performance. Conversely, the GenScore instances match the high docking power of AutoDock Vina and significantly outperform it in the screening task, achieving exceptionally high enrichment factors across the CASF2016 targets.

In the **scoring power** task (Fig. 1c), the performance of both models increases sharply within the first 100 epochs before plateauing at a Pearson correlation coefficient of approximately 0.8, significantly outperforming AutoDock Vina. However, the sequence-based XGBoost baseline achieves a comparable performance level, and even the simple KNN model outperforms Vina. This shows that a simple XGBoost model trained on top of language model embeddings can equal the performance of a complex, structure-based GNN approach when training on PDBbind and evaluating scoring power on CASF2016. Furthermore, the success of the memorization-based KNN model, which only performs nearest-neighbor searches within the training data, hints at the substantial advantage that a large dataset of affinity-labeled complexes can provide to high-capacity ML models capable of memorizing this dataset.

### Level 1: Removal of Encoding Bias

GNN-based SFs usually require encoding 3D protein-ligand structures into graph representations before processing them through the neural network. The ligand is encoded as a molecular graph with node and edge features representing chemical properties. Similarly, the protein is represented as a graph embedded with residue-specific features. Rather than modeling the entire protein, the representation is usually restricted to the binding pocket, specifically, those residues located within a defined radius of the ligand atoms. These ligand and protein graphs are then integrated into a combined interaction graph, and model predictions are generated based on the processing of this unified structure.

#### Introducing Bias During Pocket Extraction

A deeper analysis of graph dataset construction in GNN-based SFs reveals a structural bias that can inflate docking and screening performance. We define this phenomenon as “pocket bias”. As previously described, docking and screening tasks involve identifying a small set of near-native poses hidden among a much larger set of decoy molecules. For each benchmark target, the SF must detect and positively rank the near-native poses relative to the decoys, which deviate from the native pose to varying degrees in positioning and geometry.

Pocket bias is introduced when GNN pipelines construct interaction graphs for these native and decoy interactions using a single pocket representation. For each benchmark target, these pipelines extract **a single pocket graph** using a fixed radius around the **native** ligand and subsequently combine this pocket representation with all corresponding decoy ligand graphs while maintaining the true absolute coordinates of both. “Pocket bias” occurs here because the target protein’s binding site is geometrically defined and cropped based on the coordinates of a single, native reference ligand. The native ligand inherently becomes the geometric centroid of the resulting protein graph. Its atoms will have a very specific, statistically tight distribution of Euclidean distances to the surrounding protein nodes. Because they aren’t the reference point used to crop the pocket graph, the decoys will often occupy slightly different sub-regions of this pocket and their distance distributions to the pocket nodes appear “off-center” and skewed compared to the native ligand.

Two prominent examples of GNN-based SFs that employ this biased pocket construction pipeline are GenScore [16] and RTMScore [17]. Both models combine decoy ligand graphs with a static pocket representation extracted around the native ligand when they evaluate docking and screening power on the CASF2016 benchmark. Consequently, these models apply a different pocket construction methodology during benchmark evaluation than they do during training. The effect of pocket bias on decoy graphs becomes apparent when analyzing graphs from GenScore’s docking and screening evaluation.

#### Pocket Bias in Docking Power Evaluation

In the docking power benchmark dataset, all decoy poses contain the same native ligand molecule in different binding conformations, but with varying root-mean-square deviations (RMSD) from the native pose. In GenScore’s graph representations generated for this dataset, the shift in decoy ligand positioning relative to the pocket graph is clearly visible: With increasing deviation from the native pose, the ligand’s center of mass (COM) systematically drifts away from the pocket’s COM, which remains fixed around to the native ligand’s coordinates. This shift forces high-RMSD decoy ligands into off-center regions of the graph representation (Fig. 2).

**Figure 2:**
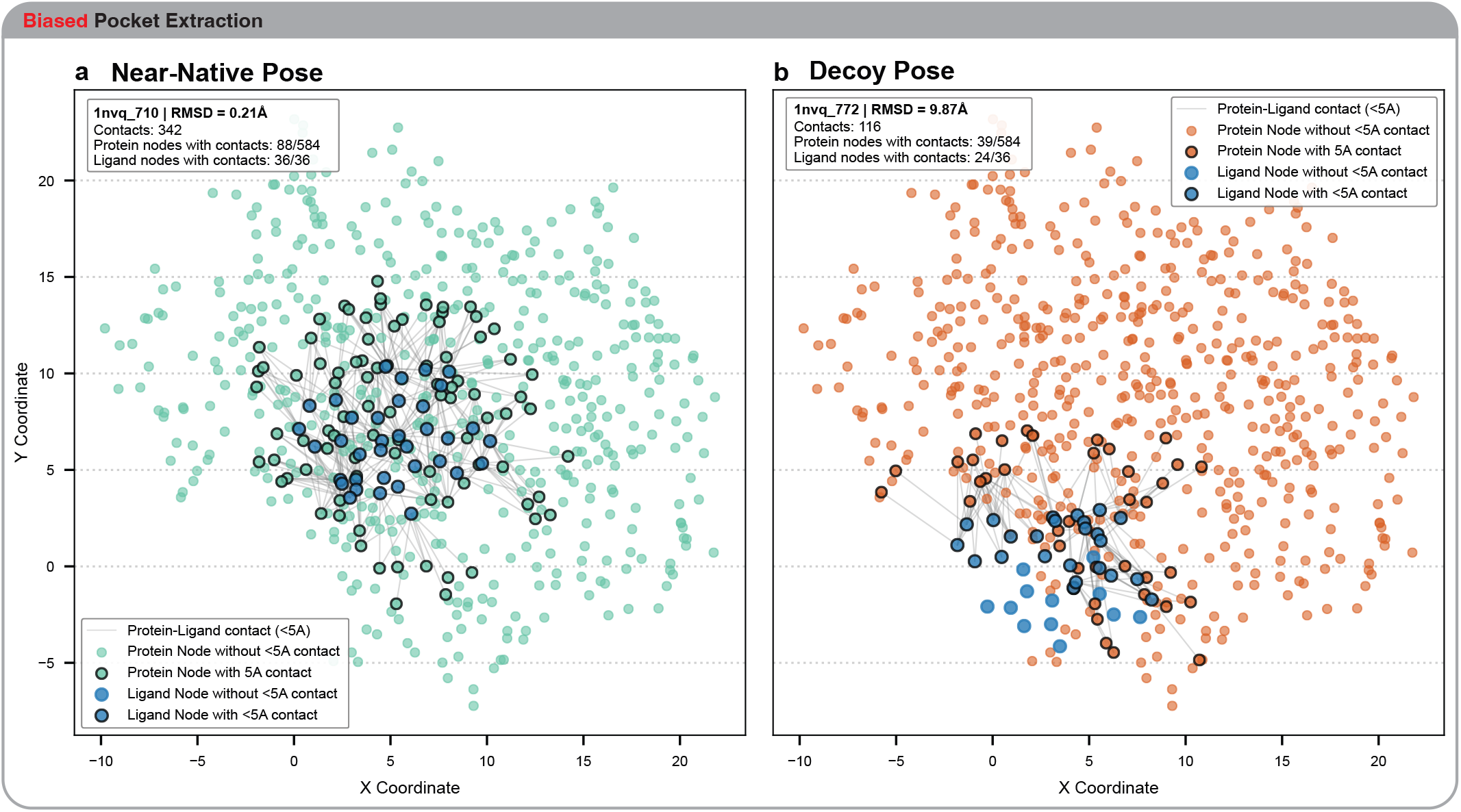
Comparison of a near-native ligand and a decoy ligand’s positioning within the pocket graph: Scatterplots of projected absolute node coordinates extracted from GenScore’s graph representations generated for the CASF2016 docking power task. The comparison highlights a) a near-native ligand and b) a high-RMSD decoy pose of the same molecule (1NVQ, UCN-01 inhibitor docked into serine-threonine kinase Chk1). With increasing RMSD, the ligand’s center of mass (COM) and the pocket’s COM drift apart, as the pocket representation remains fixed around the native reference. Consequently, ligands are located centrally in near-native graphs, while high-RMSD decoy ligands are positioned in distal regions of the graph representations. In these off-center decoy graphs, there are fewer close contacts, defined as protein and ligand nodes that are separated by less than 5 Å. For visualization purposes, coordinates are projected onto the X-Y-plane and original graph edges are omitted. Grey connections highlight “5A Contacts”, atom-residue pairs that are less than 5 Å apart in the 3D graph.

For a decoy ligand with significant deviation from the native pose, some close protein residues may have been excluded during the pocket extraction step because the native ligand was not near them. This leads to the absence of critical short-distance edges in the graph. Conversely, a decoy ligand may be embedded within a protein graph containing numerous nodes to which it is not actually proximal, but which were included due to their proximity to the native ligand. This leads to the introduction of long-distance edges into the graph of decoys, which do not exist in graphs of native or near-native ligands.

To empirically demonstrate the presence of pocket bias and the structural artifacts it introduces into decoy graphs, we computed simple structural metrics across all interaction graphs within the docking power benchmark dataset. To show how these artifacts make decoy graphs more distinguishable from near-native graphs, we subsequently implemented naive classifiers designed to separate decoy and near-native poses using these structural metrics alone.

- **Pairwise Distance Variance** Under pocket bias, decoy ligands are positioned further from the pocket graph than near-native ligands. This introduces long-range edges into the graphs and increases the variance of the pairwise distances between ligand and pocket nodes (Fig. 3a,b). A naive classifier that relies only on this variance achieves remarkably high predictive power in distinguishing near-native and decoys poses (ROC-AUC=0.665, Fig. 3c).
- **Ligand centrality:** Pocket bias shifts decoy ligands into off-center regions of the graphs. Consequently, the mean pairwise protein-ligand distance increases sharply as the decoy’s RMSD rises (Suppl. Fig. 1 a,b). A simple classifier using this mean distance achieves a notable separation of decoys from near-native structures (ROC-AUC = 0.675, Suppl. Fig. 1 c).
- **COM-distance:** Under pocket bias, the distance between the ligand and pocket COM increases drastically for high-RMSD decoy poses (Suppl. Fig. 2 a,b). Accordingly, this COM-distance entails substantial predictive power to separate near-native from decoy poses (ROC-AUC = 0.744, Suppl. Fig. 2 c).

**Figure 3:**
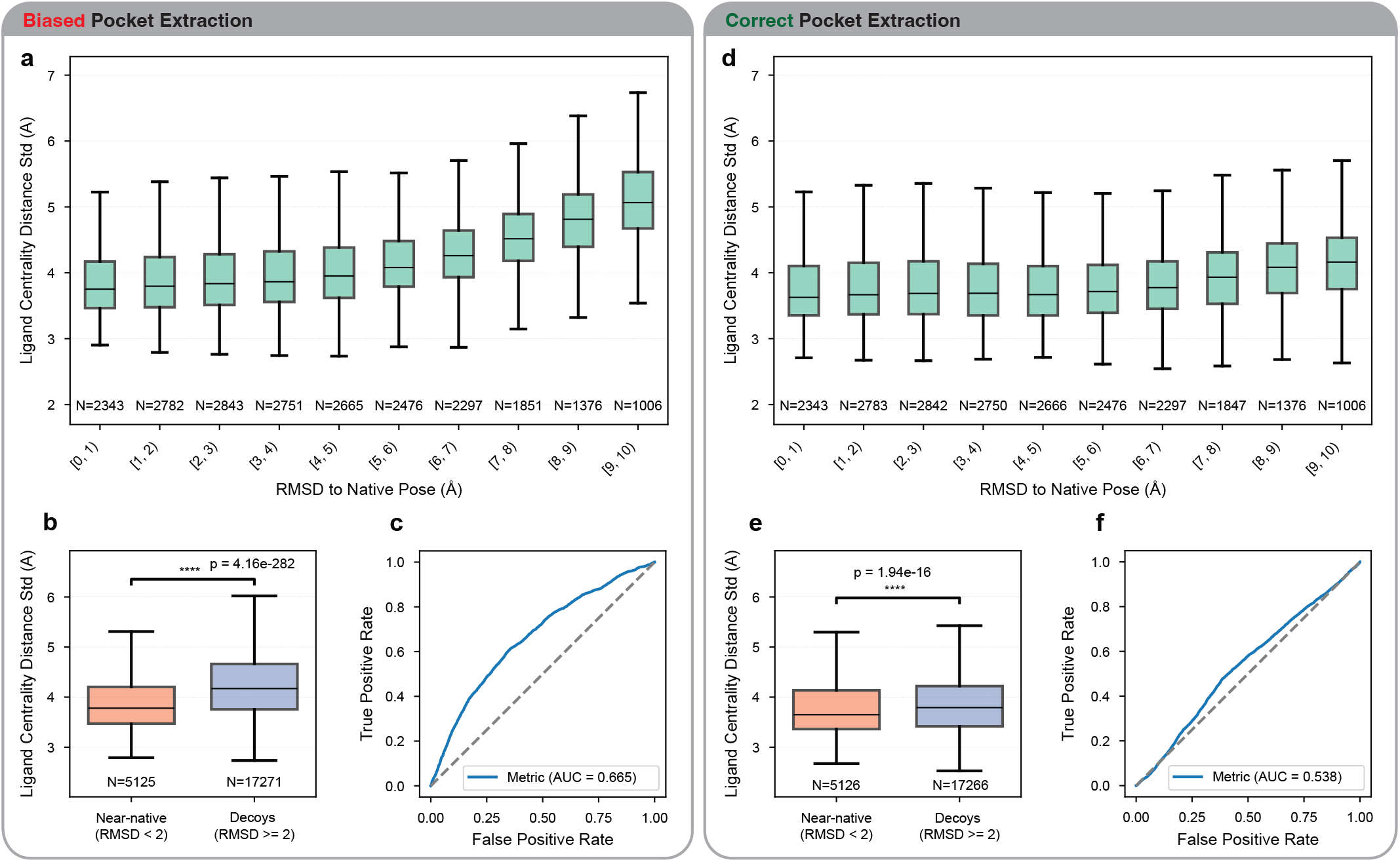
Pocket bias near-native detection through protein-ligand distance variance: Due to pocket bias, the distribution of pairwise distances between protein and ligand nodes is statistically constrained for near-native ligands but becomes significantly skewed for decoys. The decoupling of ligand positioning from pocket extraction introduces additional long-distance edges, inflating the standard deviation (SD) of these distance distributions for decoy graphs. **a, d)** Correlation between pairwise distance SD and ligand RMSD under pocket bias versus individualized pocket extraction. **b, e)** Distribution shifts in pairwise distance SD for native versus decoy graphs. **e, f)** ROC-AUC curves for a naive classifier using solely the pairwise distance SD to distinguish near-native poses from decoys. The separability is strongly increased by pocket bias. Statistical annotations include p-values computed by two-sided Mann-Whitney U tests. Boxplots represent the median (center line), 25th–75th percentiles (box), and whiskers extending to 1.5 × the interquartile range (IQR); outliers are omitted for clarity. Sample sizes (*n*) are indicated for each distribution.

#### Removing Pocket Bias – Individualized Pocket Extraction

To establish an unbiased baseline with pocket bias removed, we implemented a corrected variant of GenScore’s data processing pipeline. In this implementation, we modified the graph encoding workflow to apply pocket extraction to each binding pose individually. Instead of extracting a single protein graph around to the native ligand, this version generates a unique protein graph for every ligand pose based on its specific coordinates, mirroring the procedure used for GenScore’s training data.

When computing the same structural metrics across these corrected docking power graphs, it becomes apparent that the decoy and near-native graphs are much more structurally similar and harder to distinguish. For instance, anomalous long-range edges are eliminated from the decoy graphs, rendering the pairwise distance variances in decoy and near-native graphs more comparable (Fig. 3 d,e). As a result, the performance of the naive classifier based on this variance drops to just slightly above random chance (ROC-AUC = 0.538). Furthermore, decoy ligands are positioned more centrally within the interaction graphs, which substantially lowers the separability of native and decoy poses based on ligand centrality (ROC-AUC = 0.568, Suppl. Fig. 1 d,e,f). Finally, the center-of-mass (COM) drift is significantly reduced, reducing its classification power (ROC-AUC = 0.613; Suppl. Fig. 2 d,e,f). The systematic elevation of these ROC-AUC scores under pocket bias (Table 1) provides quantitative evidence that a baseline signal is embedded within the geometry of the graphs when using GenScore’s original pocket extraction pipeline. This represents a “geometric” data leakage and facilitates separation of decoys and near-native ligands in the CASF2016 docking power task.

**Table 1:** Simple geometric metrics achieve notable classification accuracy w ith pocket bias: ROC-AUC achieved by simple geometric metrics under biased pocket extraction versus individualized pocket extraction in the docking power benchmark. The simple metrics are directly used to classify poses into near-native and decoys. The ROC AUC score (Area Under the Receiver Operating Characteristic Curve) measures a the metric’s ability to separate the two classes across all thresholds, with scores ranging from 0 to 1. Compared to individualized pocket extraction for every pose, the pocket bias reinforces the simple geometric flags in the decoy graphs, increasing COM distances, decreasing ligand centrality and increasing the pairwise distance spread, thereby making decoys and natives more easily separable.

| Metric | Pocket Bias | Unbiased (individualized) | Improvement by pocket bias |
| --- | --- | --- | --- |
| COM distance | 0.744 | 0.613 | **+0.131** |
| Variance in protein-ligand distances | 0.665 | 0.538 | **+0.127** |
| Ligand centrality | 0.675 | 0.568 | **+0.107** |

#### Pocket Bias in Screening Power Evaluation

The screening power task, where native ligands must be identified from a chemically diverse ensemble of decoys, is inherently more demanding than the docking power task because it requires both accurate pose recognition and reliable estimation of binding strength to correctly rank active compounds above decoys. Pocket bias remains present: Because the protein graph is restricted to residues around the native ligand, the representation is artificially sculpted to fit the native ligand’s shape. If a decoy deviates from the native ligand in positioning and geometry, close protein residues may be missing from the graph because they were far from the native reference. Additionally, the decoy may be embedded within a protein graph containing many nodes to which it is not proximal. This results in the omission of short-range edges and the inclusion of extraneous long-range edges within the graphs. In some cases, this decoupling of pocket extraction from the actual ligand position generates graph representations with heavily off-center ligand graphs that would look entirely different if protein pockets were extracted around each ligand individually (Fig. 4).

**Figure 4:**
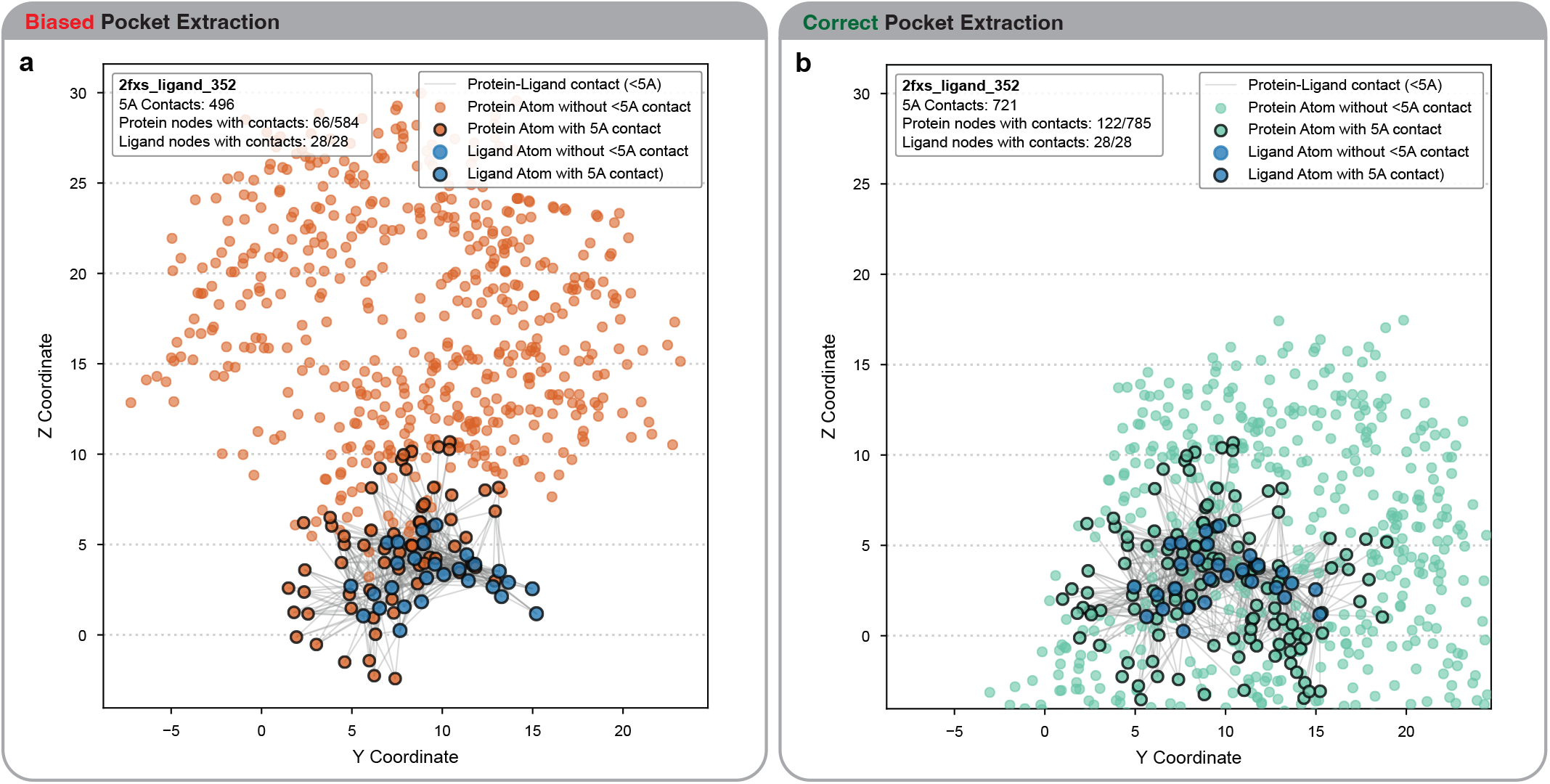
Screening Power Decoy under Biased and Individualized Pocket Extraction: Scatterplots of projected absolute node coordinates extracted from GenScore’s graph representations generated for the CASF2016 screening power task. The comparison shows spatial node distributions for the same non-binding decoy structure (HSP90 inhibitor radamide docked into Chk1 serine-threonine kinase 1NVQ) obtained with a) GenScore’s biased pocket extraction and b) individualized pocket extraction. Under pocket bias, the pocket graph extracted around the native ligand of the serine-threonine kinase is used for all decoys docked into that target protein. Because the HSP90 inhibitor is docked into a different subregion of the protein, this rigid pocket definition pushes the ligand into an off-center position of the graph representation. When pockets are extracted individually for each decoy, the graph composition changes entirely. New protein nodes are introduced, while nodes that were proximal only to the native reference ligand are removed. Under this individual pocket extraction, decoy ligands are located more centrally within the graphs and more of the essential short-distance interactions to the protein are included. For visualization purposes, coordinates are projected onto the Y-Z-plane and original graph edges are omitted. Grey connections highlight “5A Contacts”, atom-residue pairs that are less than 5 Å apart in the 3D graph.

Visual inspection clearly demonstrates the presence of pocket bias within GenScore’s screening power graphs. Moreover, major size bias exists in CASF2016 screening power benchmark. In this dataset, the native ligands are systematically much larger than the accompanying decoys (see Supplementary Note 1.3). Consequently, even properly positioned native ligands tend to appear less centrally located and exhibit larger COM distances.

#### Effect of Pocket Bias on CASF2016 Performance

To quantify the impact of pocket bias on GenScore’s benchmark docking and screening performance, we generated unbiased graph representations of the CASF2016 docking and screening power benchmark datasets using individualized pocket extraction and used these “corrected” graphs to compute docking and screening power metrics. As a result, both docking and screening power increase at a slower rate during training and manifest at a lower level upon completion (Fig. 5a,b). Notably, for the docking power task, GenScore’s performance drops below the classical AutoDock Vina baseline. These results indicate that pocket bias-induced artifacts built into decoy graph representations significantly contributes to GenScore’s high benchmark performance. The scoring power metric remains unaffected by pocket bias, because the scoring benchmark lacks decoy poses that could incorporate pocket bias. Similarily, GEMS does not include pocket bias, as its graph encoding pipeline extracts pockets individually for every pose.

**Figure 5:**
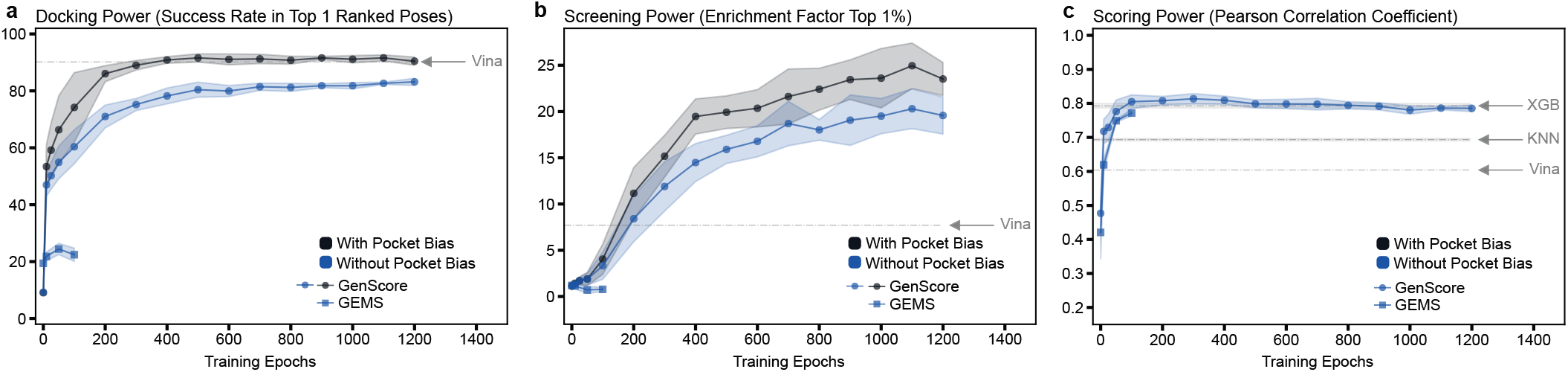
Evolution of CASF2016 performance with removed pocket bias: Comparison of **a)** docking, **b)** screening and **c)** scoring performance of GenScore, GEMS and AutoDock Vina. The scoring power plot includes two sequence-based classical machine learning models (XGBoost and *K*-Nearest-Neighbor) trained on language model embeddings. GenScore’s original docking and screening power (black) is significantly reduced without pocket bias (blue, individualized pocket extraction). In all plots, the x-axis shows training progress (epochs), while the y-axis displays the Pearson Correlation coefficient (↑, for scoring), the success rate (↑, for docking) and the enrichment factors (↑, for screening). For machine learning-based models, each line represents the mean performance across five cross-validation (CV) folds, with shaded uncertainty regions (±1 standard deviation) indicating the variability in performance across the five training runs. To maintain statistical rigor when different model instances terminated at different epoch numbers, only epochs with at least three active CV models contributing performance data are plotted.

### Level 2: Removal of Structural Data Leakage

In the context of ML-SFs, performance reporting is often significantly influenced by structural data leakage. Data leakage occurs when a ML model is evaluated on targets that share high structural similarity with targets already present in its training set. As established by previous research [15, 23, 32–37], significant data leakage exists between the PDBbind database and the CASF2016 benchmark dataset. Consequently, the benchmark metrics of high-capacity ML models are usually substantially elevated when they are trained on PDBbind and evaluated on CASF2016.

To account for this artifact, we substituted the standard PDBbind training set with a structurally deduplicated version of PDBbind, named CleanSplit, which was published alongside the GEMS model [15]. We subsequently retrained all ML models on this 5% smaller training dataset and tracked their CASF2016 benchmark performance (Fig. 6). As a result, the scoring performance of all machine learning models drops substantially, including the sequence-based XGBoost and KNN baselines. GenScore’s docking and screening power increases more slowly during the training process and ends up substantially lower than when the model is trained on the standard PDBbind dataset. GEMS’ performance on the docking and screening tasks remains consistently low, showing no further decline.

**Figure 6:**
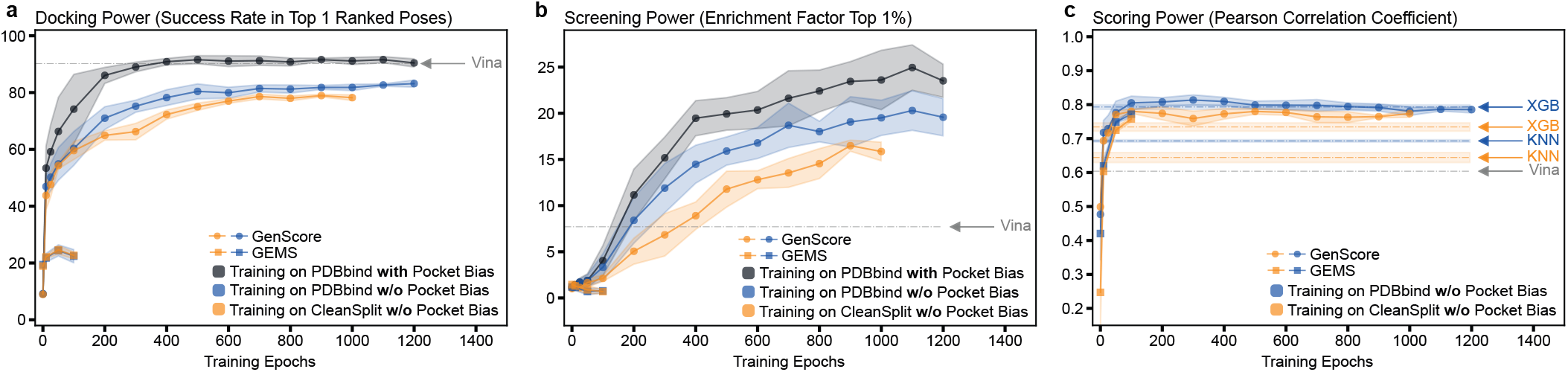
Evolution of CASF2016 performance with removed pocket bias and data leakage: Comparison of **a)** docking, **b)** screening and **c)** scoring performance of GenScore, GEMS and AutoDock Vina. The scoring power plot includes two sequence-based classical machine learning models (XGBoost and *K*-Nearest-Neighbor) trained on language model embeddings. GenScore’s original docking and screening performance (black) is significantly reduced after removal of pocket bias (blue). Across all machine-learning models, training on PDBbind CleanSplit reduces train-test data leakage and leads to a further reduction of performance (orange curves). In all plots, the x-axis shows training progress (epochs), while the y-axis displays the Pearson Correlation coefficient (↑, for scoring), the success rate (↑, for docking) and the enrichment factors (↑, for screening). For machine learning-based models, each line represents the mean performance across five cross-validation (CV) folds, with shaded uncertainty regions (±1 standard deviation) indicating the variability in performance across the five training runs. To maintain statistical rigor when different model instances terminated at different epoch numbers, only epochs with at least three active CV models contributing performance data are plotted.

In contrast, the AutoDock Vina baseline remains unaffected, as it is not a data-driven ML model that can exploit data leakage, but a physics-based empirical scoring function that assigns scores using a fixed empirical function based on physically motivated interaction terms. This leakage-free evaluation further narrows the performance gap between the GenScore and AutoDock Vina, with GenScore falling even further behind the classical baseline in the docking power task.

### Level 3: Evaluation Under Distribution Shift (OOD Datasets)

For usability in a real-world VS setting, a ML-SF should generalize effectively to target proteins and ligand scaffolds that are underrepresented within its training data. For this reason, testing a machine learning model on OOD data, which is structurally distinct from the training distribution, is essential. This validation ensures that the model can make reliable predictions when processing novel therapeutic molecules or emergent targets [21]. Ultimately, OOD testing reveals whether a model has learned to extrapolate based on an understanding of fundamental molecular interactions, rather than merely interpolating between familiar components within a well-characterized training manifold.

To perform this test of generalization, we trained seven instances of each ML-SF, each with a specific protein family explicitly excluded from the training datasets, which then served as an external test sets. These protein families were selected to represent a diverse range of protein folds, encompassing all instances of alpha-carbonic anhydrases, HIV-proteases, serine-threonine protein kinases, estrogen receptors, urokinase-type plasminogen activators, HSP82 chaperones and a cluster of transporter proteins. When tested on these completely unseen protein folds, all ML-based models exhibited severely degraded performance across the scoring, docking, screening power (Fig. 7 and Fig. 8). Their performance on the full CASF benchmark dataset remains high, confirming that their success relies on encountering familiar data distributions.

**Figure 7:**
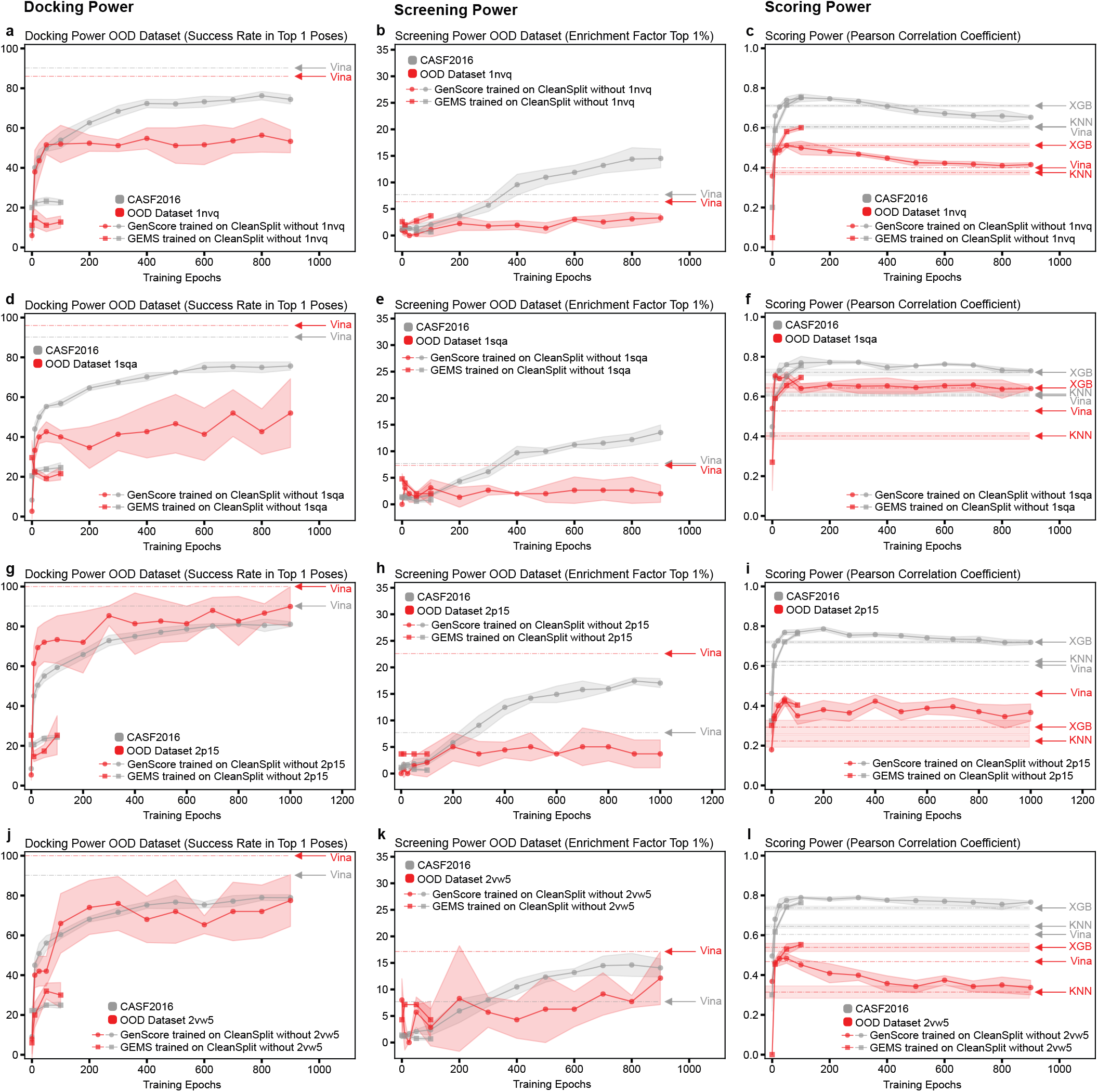
Evolution of docking, screening, and scoring performance on OOD datasets – Part 1: Comparison of **a, d, g, j)** docking, **b, e, h, k)** screening, and **c, f, i, l)** scoring performance on the complete decoy datasets of the CASF2016 benchmark (gray) and the out-of-distribution (OOD) cluster subsets (red) for GenScore, GEMS, AutoDock Vina, and two sequence-based scoring models (XGBoost and *K*-Nearest Neighbor) trained on language model embeddings. The OOD cluster datasets consist of specific protein families excluded individually from the training dataset to serve as additional OOD test datasets, namely serine–threonine protein kinases (1nvq), urokinase-type plasminogen activators (1sqa), estrogen receptors (2p15) and HSP82 chaperones (2vw5). The x-axis indicates training progress (epochs), while the y-axis displays the Pearson correlation coefficient (↑, for scoring), the success rate (↑, for docking), and the enrichment factor (↑, for screening). For machine learning-based models, each line represents the mean performance across five cross-validation (CV) folds, with shaded regions indicating the standard deviation (±1 SD). To maintain statistical rigor when model instances terminated at different epochs, only training epochs with at least three active CV models contributing performance data are plotted.

**Figure 8:**
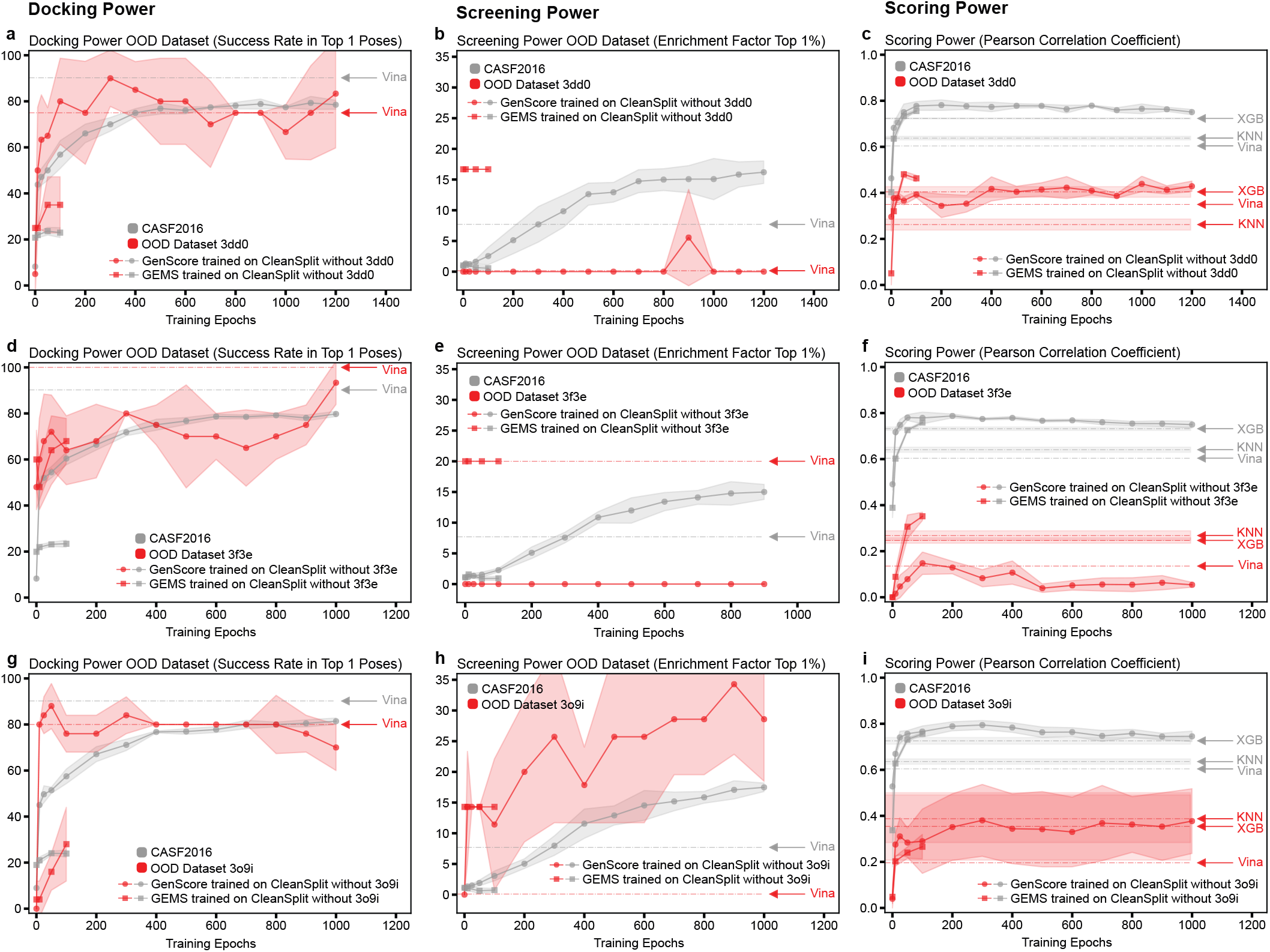
Evolution of docking, screening, and scoring performance on OOD datasets – Part 2: Comparison of **a, d, g)** docking, **b, e, h)** screening, and **c, f, i)** scoring performance on the complete decoy datasets of the CASF2016 benchmark (gray) and the out-of-distribution (OOD) cluster subsets (red) for GenScore, GEMS, AutoDock Vina, and two sequence-based scoring models (XGBoost and *K*-Nearest Neighbor) trained on language model embeddings. The OOD cluster datasets consist of specific protein families excluded individually from the training dataset to serve as additional OOD test datasets, namely alpha-carbonic anhydrases (3dd0), HIV-proteases (3o9i) and a cluster of transporter proteins (3f3e). The x-axis indicates training progress (epochs), while the y-axis displays the Pearson correlation coefficient (↑, for scoring), the success rate (↑, for docking), and the enrichment factor (↑, for screening). For machine learning-based models, each line represents the mean performance across five cross-validation (CV) folds, with shaded regions indicating the standard deviation (±1 SD). To maintain statistical rigor when model instances terminated at different epochs, only training epochs with at least three active CV models contributing performance data are plotted.

In the **scoring power** task (Fig. 7c,f,i,l and Fig. 8c,f,i), all evaluated models showed a marked decrease in performance on the OOD datasets compared to the metrics achieved on the standard CASF2016 benchmark. GEMS proved to be the most robust architecture for scoring, outperforming all other models across the majority of the OOD datasets (Tab. 2). This resilience aligns with its design objective, as GEMS was explicitly optimized to maximize scoring generalization. The sequence-based XGBoost baseline achieved the second-highest Pearson correlation coefficients, outperforming GenScore on several OOD datasets. The classical scoring function AutoDock Vina demonstrated very limited scoring performance, ranking behind GEMS, XGBoost, and GenScore. Finally, the instance-based KNN model expectedly showed the poorest scoring power. This also confirms that training data memorization is of little value when scoring these strict OOD datasets.

**Table 2:** Performance ranking of scoring functions across seven out-of-distribution (OOD) datasets, including Autodock Vina (Vina), GenScore, GEMS and the two sequence-based baseline models *K*-nearest-neighbor (KNN) and XGBoost (XGB). Cell values represent the relative rank of each model per dataset (1 to 5, where 1 indicates the best performance). The bottom row shows the mean rank across all datasets, representing average model robustness to distribution shifts.

| OOD Dataset | Docking Power |  |  | Screening Power |  |  | Scoring Power |  |  |  |  |
| --- | --- | --- | --- | --- | --- | --- | --- | --- | --- | --- | --- |
|  | Vina | GenScore | GEMS | Vina | GenScore | GEMS | GEMS | XGBoost | GenScore | Vina | KNN |
| 1nvq | 1 | 2 | 3 | 1 | 3 | 2 | 1 | 2 | 3 | 4 | 5 |
| 1sqa | 1 | 2 | 3 | 1 | 2 | 2 | 1 | 3 | 2 | 4 | 5 |
| 2p15 | 1 | 2 | 3 | 1 | 2 | 3 | 2 | 4 | 3 | 1 | 5 |
| 2vw5 | 1 | 2 | 3 | 1 | 2 | 3 | 1 | 2 | 4 | 3 | 5 |
| 3dd0 | 2 | 1 | 3 | 2 | 2 | 1 | 1 | 3 | 2 | 4 | 5 |
| 3f3e | 1 | 2 | 3 | 1 | 2 | 1 | 1 | 3 | 5 | 4 | 2 |
| 3o9i | 1 | 2 | 3 | 3 | 1 | 2 | 4 | 3 | 2 | 5 | 1 |
| **Average Rank** | **1.14** | **1.86** | **3.00** | **1.43** | **2.00** | **2.00** | **1.57** | **2.86** | **3.00** | **3.57** | **4.00** |

**Table 3.** Hyperparameter search space for the K-Nearest Neighbors (KNN) regressor.

| Parameter | Distribution/Options | Range/Values |
| --- | --- | --- |
| n_neighbors | Uniform Integer | [3, 30] |
| weights | Categorical | uniform, distance |
| metric | Categorical | euclidean, manhattan, minkowski |
| p | Uniform Integer | [1, 5] |

**Table 4.** Hyperparameter search space for the XGBoost regressor. Note that uniform(a, b) denotes a distribution starting at *a* with width *b*.

| Parameter | Distribution | Range |
| --- | --- | --- |
| n_estimators | Uniform Integer | [300, 1200] |
| learning_rate | Log-Uniform | $[10^{-3}, 10^{-1}]$ |
| max_depth | Uniform Integer | [3, 8] |
| subsample | Uniform | [0.7, 1.0] |
| colsample_bytree | Uniform | [0.7, 1.0] |
| reg_alpha | Log-Uniform | $[10^{-3}, 10]$ |
| reg_lambda | Log-Uniform | [0.1, 50] |

In the **docking power** task (Fig. 7a,d,g,j and Fig. 8a,d,g), AutoDock Vina dominated the evaluation, outperforming both GenScore and GEMS across all but one OOD dataset (Tab. 2). GenScore ranked second, exhibiting decent docking power generalization by maintaining success rates comparable to those achieved on the full CASF2016 docking benchmark. In contrast, GEMS demonstrated poor performance, showing very limited docking utility.

In the **screening power** task (Fig. 7b,e,h,k and Fig. 8b,e,h), AutoDock Vina outperformed the ML-SFs on five out of seven OOD datasets (Tab. 2), often maintaining or exceeding its performance metrics from the full CASF2016 screening benchmark. In stark contrast, Vina registered enrichment factors of zero for both the alpha-carbonic anhydrases and HIV proteases. Interestingly, GenScore exhibited sharp reductions in its enrichment factors across all but one OOD dataset, nearly closing the performance gap to the underperforming GEMS model, which was optimized exclusively for the scoring task. This suggest that GenScore’s claimed superior benchmark screening performance is entirely based on a combination of pocket bias and encountering in-distribution data during evaluation.

## Discussion

Overall, our results indicate that the apparent performance advantages of GNN-based SFs over classical SFs are highly sensitive to evaluation design. Across CASF2016 scoring, docking, and screening tasks, GenScore and GEMS achieve strong headline performance under standard evaluation, but we find that these metrics are partly driven by systematic biases. When these sources of bias are progressively removed, the performance of these models drops substantially, in several settings approaching or falling below the classical AutoDock Vina baseline. Removing train-test overlap via deduplicated training data lowers performance for all tested ML-based models. These findings reinforce a key point: High performance of ML models often reflects the presence of near-neighbors in the training distribution. As a consequence, comparisons to classical SFs that do not explicitly control structural overlap represent an unfair comparison. The most stringent evaluation condition involves testing on OOD data, which we believe provides the most accurate reflection of a model’s utility for prospective VS campaigns on novel targets. In this OOD evaluation, the classical SF AutoDock Vina proved superior across both docking and screening tasks. Only in the scoring of OOD data do the ML models maintain an advantage over Vina, led predominantly by the relatively simple GEMS model. This superiority of Vina suggests higher stability under distribution shift, consistent with its reliance on more general physicochemical priors. For drug discovery, where novelty is often the point, robustness under distribution shift is arguably more important than in-distribution benchmark scores.

### Data Processing Can Introduce Bias

A central finding of this work is that seemingly minor data processing decisions can introduce strong geometric data leakage. The state-of-the-art ML-SFs GenScore and RTMScore incorporate a data encoding pipeline that extracts pocket residues around the coordinates of the native ligands and combines the resulting graph with all decoy structures. While this strategy might seem logical considering a drug discovery setting with a fixed known target pocket, it clearly introduces a bias in the context of benchmarking against other models on CASF2016. Pocket construction anchored to the native ligand embeds artifacts into the graphs, which make decoy ligands locate in “off-center” region of the pocket graphs and lead to the omission of crucial short-distance interactions. With this bias, the graph encoding process employed for the CASF2016 docking and screening decoys systematically differs from the procedure that generated GenScore’s training data. Correcting this encoding, reverting it to the standard pocket extraction used for generating the training graphs, substantially reduces benchmark docking and screening scores.

### Docking and Screening Performance Depend on OOD Failure Mode

Both GEMS and GenScore are trained on the general set of the PDBbind database, which comprises only native binders. Consequently, these models are trained only on positives, yet are asked to reject a large number of non-binding decoys during docking and screening power evaluations. Theoretically, at the current scale of labeled datasets, a model trained on only positive data should be incapable of learning the decision boundaries necessary to distinguish decoys from native poses. In line with this expectation, GEMS exhibits poor docking and screening performance. However, even after removing pocket bias and structural data leakage, GenScore maintains considerable performance under the same training constraints, especially in the docking power task. Here, we critically discuss the aspects of GenScore’s model architecture that account for its superior performance on CASF2016.

PDBbind structures come from crystallization experiments, capturing the experimentally validated thermodynamic energy minima of bound complexes. In contrast, the CASF2016 decoy sets consist of computationally generated non-binding poses that are deliberately engineered to be suboptimal. Given these different origins, it is reasonable to assume that systematic structural differences exist between the datasets, representing a significant distribution shift. Consequently, these artificial decoy poses represent unfamiliar OOD inputs for models trained only on native crystal structures. As ML models generally show lower performance on data that is OOD with respect to their training data, both GEMS and GenScore are expected to yield higher prediction errors for the artificial decoy poses.

For high performance in the CASF2016 docking and screening tasks, **it is absolutely essential that a SF assigns low scores to decoy poses**. Under the assumption that decoys represent OOD inputs, performance in these tasks depends critically on the specific failure mode of the model when encountering such data. If a model penalizes OOD inputs with low scores, its docking and screening power increases drastically. Conversely, if a model assigns high scores or simply maps OOD inputs to arbitrary values, performance diminishes sharply.

GenScore incorporates an architectural mechanism that systematically causes OOD inputs to receive low scores. Instead of directly predicting an affinity score, its Mixture Density Network (MDN) predicts the distances between all atom–residue pairs in the complex. It then evaluates how well the predicted distances match the true distances. For each pair, a log-probability is calculated, representing the likelihood of the true distance occurring under the predicted distance probability distribution. The final score is calculated as the sum of these individual log-probabilities across all atom–residue pairs. When presented with a decoy that is OOD with respect to its training data, GenScore’s distance predictions likely become inaccurate, leading to low probabilities and a correspondingly low aggregate score. Conversely, for familiar native structures, the predicted distances are likely more accurate and align more closely with the true distances, yielding high probabilities and elevated scores. Consequently, scores for OOD inputs are effectively forced to low numbers by design, increasing GenScore’s benchmark docking and screening power. In contrast, GEMS is trained on the scoring task, performing direct regression to affinity scores. It has no mechanism to detect and penalize OOD inputs. When evaluating a decoy that is OOD, GEMS likely predicts an arbitrary value from the training label distribution rather than assigning a low score.

Ultimately, the divergent performance of GEMS and GenScore on the docking and screening benchmark can be partly attributed to how they score OOD structures. While both models likely generate erroneous predictions for decoys, GEMS outputs these erroneous scores, whereas GenScore systematically maps them to a low number.

### Interplay of MDN and Pocket Bias Inflates Performance

In the original version of GenScore with pocket bias, docking and screening metrics are strongly inflated by the interplay of pocket bias and MDN. While the graphs of near-native decoys are less affected by pocket bias and remain relatively similar to the training data, pocket bias introduces structural artifacts into the graphs of high-RMSD decoys, such as off-center ligand positioning and long-distance protein–ligand edges. This further alienates the decoy poses and makes them even more OOD with respect to the training data. As an example, when processing high-RMSD decoys, the model encounters graphs full of pocket-bias-induced long-distance edges. As these do not exist in the training data, the prediction errors for such decoys will be large, which drives the output scores even lower.

In addition to making decoy graphs more OOD relative to the training data, pocket bias also lowers the output scores for decoys in a more direct way: It leads to the omission of crucial short-distance edges from decoy graphs. After predicting distances for all atom–residue pairs and computing a log-probability for each, GenScore restricts the final summation to atom–residue pairs that are separated by less than 5 Å. Because the decoy graphs generated under pocket bias contain fewer of these short-distance edges, there are fewer probabilities to sum up. This artificially depresses the output scores for decoy poses, thereby inflating the benchmark docking and screening power further.

### Size Bias in CASF2016 Benchmark Inflates Screening Performance Metrics

A substantial size bias exists within the CASF2016 screening power benchmark (Suppl. Note 1.3), where the target ligands designated for identification (Top-1 ligands) are significantly larger than the accompanying decoy ligands (Suppl. Fig. 3). This bias in the CASF2016 benchmark design artificially enhances the screening power metrics of models that positively reward molecular size. GenScore exploits this benchmark bias through its scoring mechanism, in which the final score is calculated as the sum of log-probabilities across all atom–residue pairs. Because the Top-1 ligands are significantly larger, they generally form more contacts within the 5 Å threshold than the decoys. Consequently, there are more probabilities to sum up in the final summation. Due to this hard-coded size reward, the Top-1 ligands receive systematically higher scores than the decoy ligands, simply due to their atom counts. It is therefore expected that screening performance metrics would decrease if models were evaluated on a benchmark with Top-1 ligands and their corresponding decoys sharing a similar size distribution.

### Conclusions and Considerations for Future Development of ML-SFs

The sharp performance declines observed across ML-based SFs upon the removal of target-proximal data from their training distributions suggest that these models, while acting as strong interpolators, struggle to extrapolate to unfamiliar structures. While the scoring power results of GEMS demonstrate that smaller, highly regularized models can achieve superior extrapolation, GenScore appears overfitted to the training data, yielding a model that performs exceptionally well on in-distribution data but fails when confronted with OOD targets. In contrast, the classical SF AutoDock Vina proves significantly more robust under distribution shifts.

To provide genuine value in prospective VS, future ML-SFs must aim to incorporate stronger physical priors that facilitate extrapolation to novel targets. In this context, GenScore’s architecture, particularly its use of Mixture Density Networks [42], represents an interesting approach related to pose prediction, requiring models to learn the probability distribution of the distance between each ligand-protein atom pair. Moving forward, the integration of more physical constraints could be achieved through delta-learning, an approach where a neural network is trained to predict the residual error of a physically bounded baseline scoring function rather than predicting absolute binding affinities from scratch [43].

Beyond architectural improvements, model robustness is inextricably tied to data quality and rigorous benchmark design. There is a critical need for more blinded benchmark datasets specifically designed to evaluate ML-based SFs [21]. Before choosing a benchmark, researchers must verify that it is suitable for their specific method and confirm that a dataset’s unbiasing strategy works for their architecture [25]. Future ML-SF development should transition from standard training datasets like PDBbind v.2020 to open-source, optimized datasets such as HiQBind [44], which removes problematic complexes and provides refined ligand, protein, and hydrogen placements. Furthermore, capturing the physical reality of protein-ligand binding requires modeling explicit water molecules, as water-mediated interactions are non-negligible and difficult for models to learn implicitly.

To ensure models are learning true interaction physics, complex ML methods must be tested against simple approaches, such as classical ML models or nearest-neighbor similarity searches, to prove the complex architecture is actually necessary for achieving the given results. Detecting noncausal bias is equally critical and requires applying negative controls or sanity tests, such as systematically removing the protein structure entirely from the input to confirm that the model’s predictive performance drops as expected [25]. These tests can be facilitated by ToolBoxSF [24], an easy-to-use platform for interrogating scoring function performance.

Finally, we advocate for more rigorous evaluation paradigms: Identical data processing and graph encoding strategies used during training must be strictly maintained throughout benchmark evaluation. Deviating from this consistency risks introducing structural bias and producing artificially inflated metrics, as demonstrated by the pocket bias observed in GenScore. Furthermore, future ML-SFs should be optimized and evaluated using cross-validation rather than single data splits, and final testing protocols must explicitly account for structural overlap between the training and evaluation sets. As an ultimate proof of utility, models must demonstrate their viability for drug discovery by proving their generalizability against OOD targets that accurately reflect real-world deployment scenarios [21].

## 1 Methods

### 1.1 Evaluation Methodology and Pipeline Standardization

To rigorously assess the intrinsic algorithmic superiority of the evaluated architecture, we eliminate confounding variables introduced by differing evaluation protocols. Consequently, we implemented a strictly standardized training and evaluation pipeline across all models using a consistent 5-fold cross-validation strategy on the PDBbind dataset with identical partitioning. Each model is trained on 4/5 of the training data, with 1/5 reserved for validation. We acknowledge that this represents a slight reduction in training data volume compared to some of the original baseline implementations (GenScore used 1500 validation samples). Because deep learning models often scale with data volume, this constraint may result in a marginally lower peak performance relative to the originally published state-of-the-art metrics.

However, this trade-off is both intentional and necessary for a fair comparative analysis. By constraining the data volume and holding the validation splits identical across all experiments, we ensure the following:

- Isolation of Algorithmic Variance: Any observed delta in performance between models can be confidently attributed to predictive superiority, rather than discrepancies in training data volume or lucky random seed initializations.
- Statistical Significance: A single point estimate is highly sensitive to the specific data split. Utilizing 5-fold cross-validation allows us to report the mean performance alongside standard deviation, providing a quantifiable measure of model stability and variance that single-holdout methods lack.

While the absolute metrics reported herein may reflect the slightly constrained training volume, the relative rankings and observed performance gaps provide a more robust and statistically sound evaluation of the models’ true capabilities.

### 1.2 Datasets & Splitting

The main data resource used in this work was the PDBbind (v.2020) database [30, 31], containing 19’443 protein-ligand complexes from the Protein Data Bank (PDB) with experimentally measured binding affinities. This database is split into a general set (n=14127) and a refined set (n=5316) that has been compiled based on strict curation criteria, including crystallographic structures (excluding NMR structures) with a resolution of < 2.5Å and an inhibition constant (Ki) or dissociation constant (Kd) in the range of 1pM to 10mM (pK range 2-12). For training of all models, a merged dataset containing all data from the general and the refined set was used as training data, excluding all complexes present in the Comparative Assessment of Scoring Functions (CASF) benchmark datasets [22] (versions 2013 and 2016), which served as external test datasets in this work.

#### Removal of Train-Test Data Leakage

As a deduplicated training dataset, we used the PDBbind CleanSplit dataset (N=16909) that was published alongside the GEMS model [15]. PDBbind CleanSplit is a refined training dataset variant of PDBbind designed to eliminate redundant overrepresentation and train-test data leakage into CASF2016.

#### OOD Datasets

To enable a more stringent assessment of out-of-distribution (OOD) generalization, we constructed train–test splits derived from PLINDER’s [34] pocket-level clustering, which groups PDB entries into communities based on pocket-lDDT similarity; concretely, we use the `pocket_lddt_50_community` cluster type. We anchor each split on a different CASF-2016 representative chosen to cover a diverse range of protein families (PDB IDs `1NVQ`, `1SQA`, `2P15`, `2VW5`, `3DD0`, `3F3E`, `3O9I`) and assign every complex in the anchor’s PLINDER community to the test set, with the remaining PDBbind data used for training and validation. Throughout this section we refer to each split by its anchor (e.g. the “1NVQ cluster”). This protocol ensures that the test pockets are structurally distinct from those observed during training, thereby providing a substantially more challenging evaluation of OOD generalization.

#### Dataset Splitting

A 5-fold cross-validation split was generated for each of the following training datasets:

- Dataset 1: Complete PDBbind dataset
- Dataset 2: PDBbind CleanSplit
- Dataset 3: PDBbind CleanSplit excluding the 1nvq cluster
- Dataset 4: PDBbind CleanSplit excluding the 1sqa cluster
- Dataset 5: PDBbind CleanSplit excluding the 2p15 cluster
- Dataset 6: PDBbind CleanSplit excluding the 2vw5 cluster
- Dataset 7: PDBbind CleanSplit excluding the 3dd0 cluster
- Dataset 8: PDBbind CleanSplit excluding the 3f3e cluster
- Dataset 9: PDBbind CleanSplit excluding the 3o9i cluster

For each dataset, random 5-fold splitting was performed with stratification based on affinity labels to ensure a consistent label distribution across all folds, following the GEMS implementation. This identical stratified split was then maintained across all models trained on a given dataset, including GenScore and the sequence-based classical machine learning baselines utilized in the scoring power evaluation.

### 1.3 Model Training & Evaluation

Training of GEMS and GenScore was performed according to the original instructions using the published source code and data without modifications. The only alteration to the GenScore workflow was the implementation of the custom train-validation splits. Models were trained using the provided scripts, maintaining all original hyperparameters, training configurations, and early stopping criteria. For each model variant, five independent models were trained, corresponding to the 5-fold cross-validation (CV) scheme where each fold served as the validation set once, with intermediate checkpoint saving at regular epoch intervals. Subsequently, these checkpoints were reloaded to evaluate CASF2016 scoring, docking, and screening power using the standard evaluation scripts included in the CASF2016 benchmark. At each saved epoch, mean and standard deviation of the CASF2016 performance metrics were computed across the five CV models.

#### Evaluation on OOD Datasets

To probe generalization beyond the pockets seen during training, we construct seven out-of-distribution (OOD) test splits, each holding out an entire family of structurally similar pockets rather than a random set of complexes. Reporting metrics per cluster gives a per-family readout, which is more informative than a single aggregate number across all OOD complexes. For each cluster we report three complementary power metrics, following the CASF-2016 protocols:

- **OOD scoring power**. Pearson *R* between the predicted score and −log *K*_*d*_*/K*_*i*_ across the full PLINDER community of the cluster. For VINA, complexes whose score indicates a steric clash in the crystal pose are excluded.
- **OOD docking power**. Top-1 success rate at ≤ 2 Å heavy-atom RMSD, computed on the CASF-2016 decoy poses for those cluster members that are also CASF-2016 coreset entries (4–50 complexes per cluster).
- **OOD screening power**. Forward-screening enrichment factor *EF*_1%_ on the CASF-2016 setup, restricted to those screening targets whose PDB IDs fall in the cluster’s community (1–10 targets per cluster). For each target, the 285 candidate ligands of the CASF-2016 screening benchmark are ranked by their best-pose score; we then compute *EF*_1%_ = (*n*_top_*/n*_total_)*/*0.01, where *n*_top_ is the number of true binders recovered in the top 1% of the ranked list and *n*_total_ is the total number of true binders for that target. *EF*_1%_ = 1 corresponds to random retrieval, and higher values indicate selective enrichment of true binders. We report the per-target mean across the targets in the cluster.

#### Sequence-based baselines for the scoring power task

Sequence-based K-Nearest Neighbors (KNN) and XGBoost baselines for the scoring power task were implemented using python (3.14.5) scikit-learn (v.1.8.0), xgboost (3.2.0), scipy (1.17.1), pandas (3.0.3) and numpy (2.4.6). Data preprocessing was performed with RDKit (2024.03) and language model embeddings were generated with MolFormer [40] (`ibm/MoLFormer-XL-both-10pct`) and ESM-2 [41] (`facebook/esm2_t30_150M_UR50D`), accessed through the transformers library (4.33.3). Language model embeddings were concatenated to form the final input data vectors representing the molecular complexes.

KNN and XGBoost models were trained on the same training datasets described above, using a CV training protocol across the same predefined cross-validation folds using scaled feature embeddings and pK values as targets. We performed hyperparameter optimization via `RandomizedSearchCV` with 25 iterations and *R*^2^ scoring to identify the optimal parameter configuration. To evaluate the predictive performance of the KNN and XGBoost algorithm, the five fold-specific models were then individually evaluated on all test datasets, applying the data scaler fitted on the training set. *R*^2^ and Pearson correlation coefficients were averaged across the five models, with the standard deviation serving as uncertainty estimate.

### Evaluating GenScore without Pocket Bias

Since pocket bias only impacts the evaluation of docking and screening power (where decoys are present), only the graph construction pipeline for these benchmark poses required adaptation; no changes to the model itself were necessary. For both the training dataset and the CASF-2016 scoring power benchmark, GenScore already extracts pockets individually for each protein-ligand pair. Consequently, the pocket extraction methodology used for the training data could be reused for the docking and screening power benchmarks without modification. Only minor adjustments to the `VSDataset` function were needed to invoke the existing `extract_pocket` function for each ligand individually, rather than assigning the precomputed native ligand pocket to all pairs.

### Analysing GenScore Graph Structures

Graph representation were extracted during GenScore’s forward pass by adding saving functionality to the `run_an_eval_epoch` function, which saved coordinate arrays, the true pairwise protein-ligand distance arrays, the batch assignments, and the IDs. During the following analysis, we computed ligand and protein centers of mass (COM), COM-to-COM distance, contact counts below 5Å, atom counts, contact fractions and ligand centrality metrics (statistics over all pairwise ligand-protein node distances).

To determine the statistical significance of differences between two independent groups, a two-sided Mann-Whitney U test was performed. This non-parametric test was chosen to compare differences between groups without assuming a normal distribution of the data. For data visualizations, significance levels are denoted using the following standard conventions:

- ns (not significant) for *p* ≥ 0.05
- * for *p* < 0.05
- ** for *p* < 0.01
- *** for *p* < 0.001
- **** for *p* < 0.0001

Where applicable, precise p-values are displayed directly on the graphs above brackets indicating the compared groups. In cases of extremely high significance, p-values smaller than 1e-300 are reported as “< 1e-300”. Correlations between RMSD and spatial metrics were assessed using both Pearson (parametric) and Spearman (non-parametric) correlation coefficients. Predictive power of individual metrics for distinguishing native-like from decoy poses was evaluated via receiver operating characteristic (ROC) curve analysis, with area under the curve (AUC) computed as the discrimination metric. For RMSD-stratified analysis, complexes were binned into 1 Ångström intervals, with bins containing fewer than 1% of total data points excluded to avoid spurious statistical conclusions from sparse bins.

The absolute coordinates are used to generate comparative 2D visualizations of protein-ligand spatial relationships. For each comparison of near-native and decoys, the pipeline automatically selects the best 2D projection plane (XY, YZ, or XZ) by maximizing the separation between ligand point clouds of the two structures. All protein and ligand atoms are plotted with distinct visual differentiation between atoms involved in < 5 Å contacts versus those not in contact, with gray connection lines between interacting protein-ligand atom pairs.

### Scoring with Autodock Vina

We include AutoDock Vina v1.2.7 as a non-learned baseline. Each protein–ligand complex from PDBbind v2020 is preprocessed by adding hydrogens, assigning Gasteiger partial charges, and converting both receptor and ligand to the PDBQT format expected by Vina; the receptor is built from the pocket region. A cubic search box centered on the crystallographic ligand is then passed to Vina in `score_only` mode, which evaluates the empirical Vina scoring function on the crystal pose without re-docking.

We verify the implementation on the CASF-2016 core set and obtain Pearson *R* = 0.601 against the reported *R* = 0.604 [22]. We then apply the same protocol to the OOD clusters. For the docking and screening power evaluations we use the per-pose Vina scores shipped with the CASF-2016 benchmark package directly.

For the `3dd0` cluster, only 7 of 485 community members could be prepared from PDBbind v2020; on this cluster only, we therefore use the 2024 re-curation of PDBbind [30, 31] for scoring power, which restores 466 of the missing entries. All other clusters use PDBbind v2020.

## Data Availability

All protein–ligand complex structures and binding affinity data used in this study were obtained from the PDBbind database version 2020 [30, 31], which is publicly accessible at http://www.pdbbind.org.cn. For fast reproduction of our results, we provide PyTorch datasets of precomputed graphs for training and evaluating GenScore (https://doi.org/10.5281/zenodo.21510319) and GEMS (https://doi.org/10.5281/zenodo.14260170). The data generated in this study are freely available. The data splits used and the performance metrics of GenScore, GEMS and the classical machine learning baselines on CASF2016 and the OOD datasets can be accessed through GitHub (https://github.com/DavidGraber/PocketBias). Precomputed language model embeddings and the source data underlying the analysis of GenScore graph representations (with and without Pocket Bias) are accessible through Zenodo (https://doi.org/10.5281/zenodo.21510319).

## Code Availability

The code generated in this study is freely available on GitHub. All scripts for training, inference and evaluation of GenScore models, as well as analysis of GenScore graph representations, can be accessed through https://github.com/DavidGraber/PocketBias. This repository builds upon and modifies the original GenScore repository by sc8668, which can be accessed at https://github.com/sc8668/GenScore. The code for training and evaluating GEMS can be accessed through https://github.com/camlab-ethz/GEMS. The scripts for training and evaluating the classical machine learning baselines are accessible at https://github.com/DavidGraber/PDBbind ML baselines. All code is distributed under the MIT License.

## Author Contributions

D.G., J.K. designed and carried out the experiments

D.G. analysed the data and prepared visualizations with feedback from J.K., P.S., B.K. and S.M.

D.G., J.K, wrote the manuscript with feedback from P.S., B.K. and S.M.

R.N. trained sequence-based KNN and XGBoost baselines

The project was supervised by S.M.

## Acknowledgments

We would like to thank Prof. Dr. Rebecca Buller for her valuable feedback on this research project.

