## Supplementary Information for "How Bias Shapes the Leaderboard: Scoring Function Performance Under Scrutiny"

### Supplementary Figure 1: Docking Power Pocket Bias - Ligand Centrality

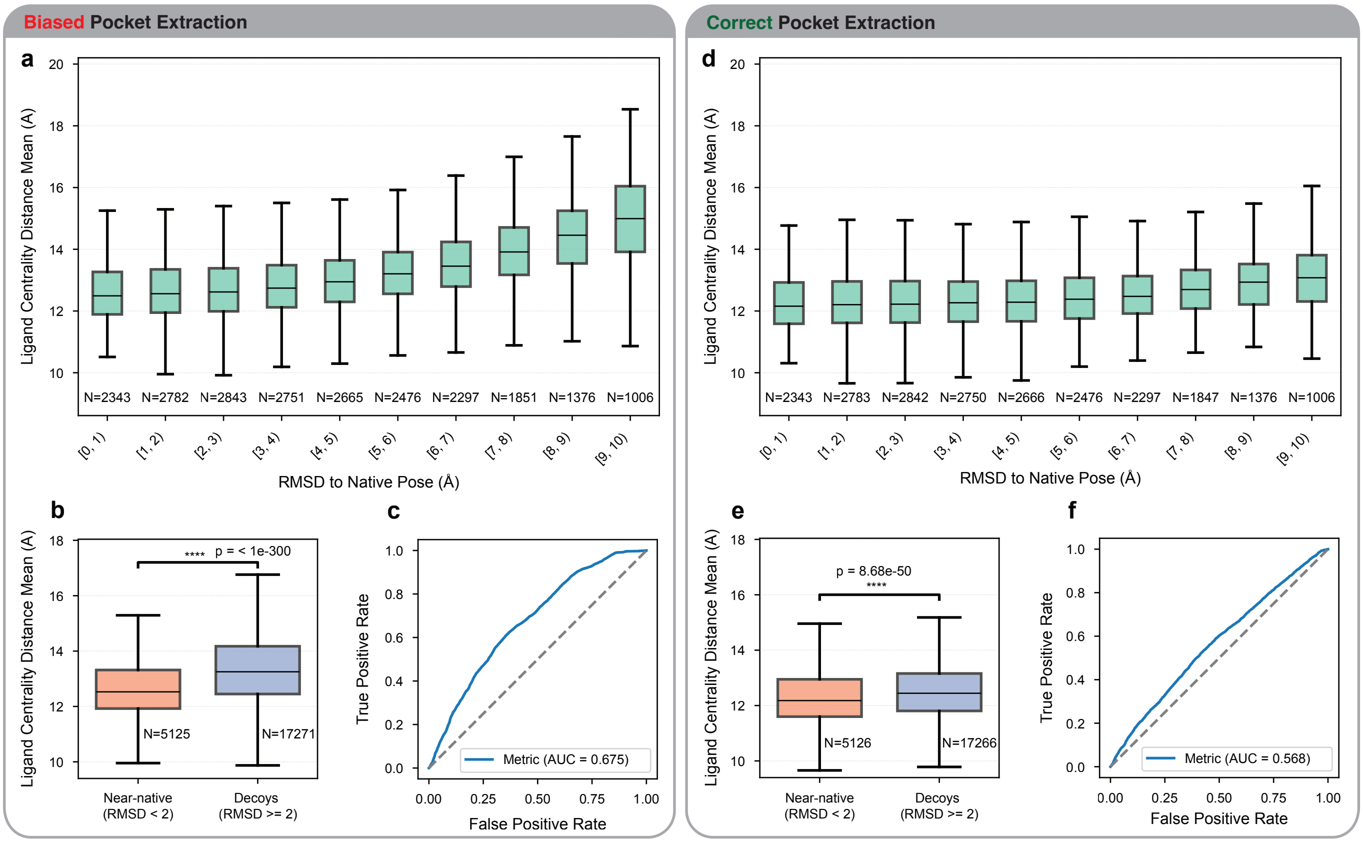


**Supplementary Figure 1: Pocket bias enables near-native detection through ligand centrality:** Due to the pocket bias, decoy ligands are shifted into heavily off-centered locations in the protein-ligand graphs. This increases the mean pairwise distance between ligand and protein nodes in these representations. **a, d)** Correlation between mean pairwise distance and ligand RMSD under pocket bias versus individualized pocket extraction. **b, e)** Distribution shifts in mean pairwise distance for near-native versus decoy graphs. **c, f)** ROC-AUC curves for a naive classifier using solely the mean pairwise distance to distinguish near-native poses from decoys. The observed separability is strongly enhanced by pocket bias. Statistical annotations include p-values computed by two-sided Mann-Whitney U tests. Boxplots represent the median (center line), 25th–75th percentiles (box), and whiskers extending to 1.5 $\times$ the interquartile range (IQR). Outliers are omitted for clarity. Sample sizes ($n$) are indicated for each distribution.

### Supplementary Figure 2: Docking Power Pocket Bias - Center-of-Mass Drift

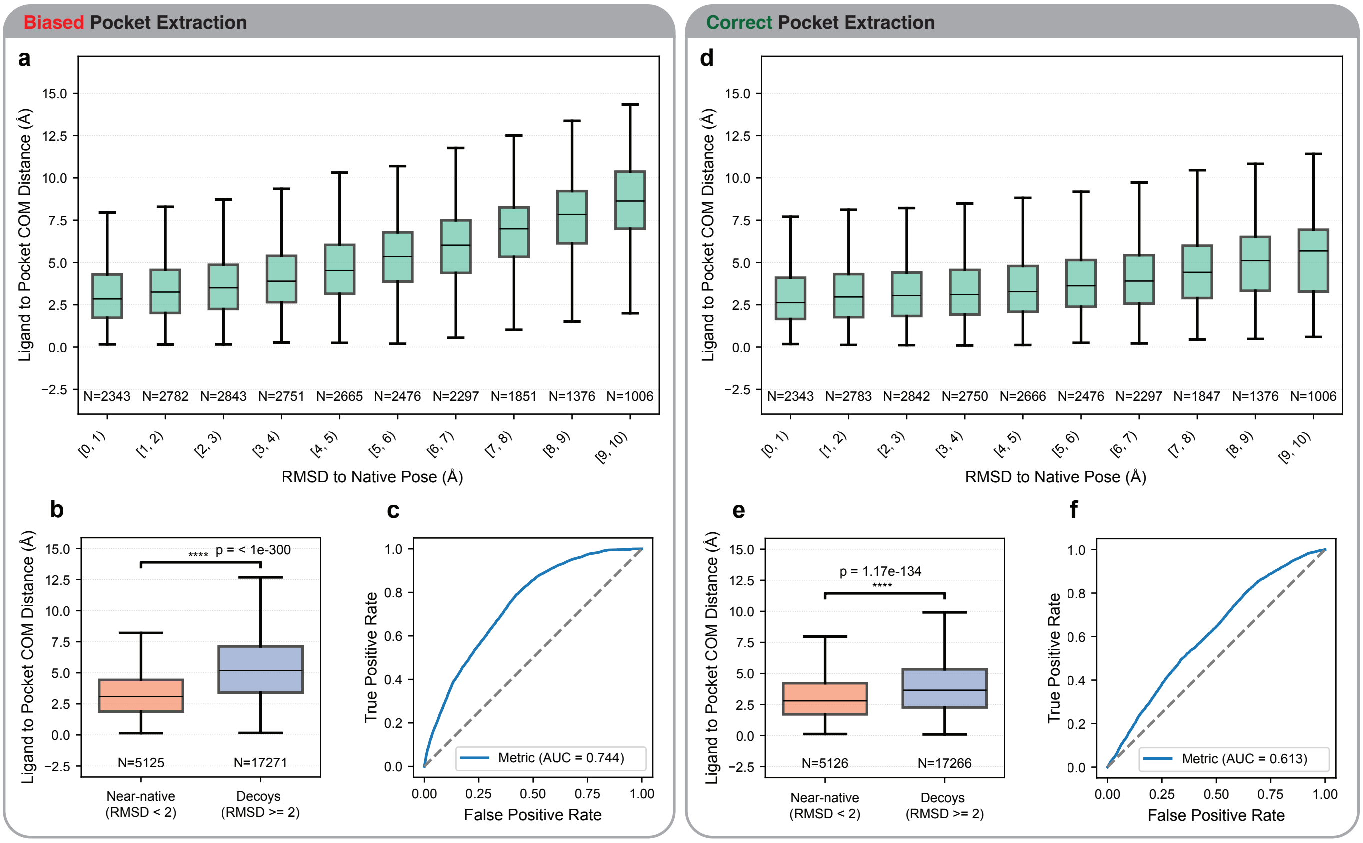


**Supplementary Figure 2: Pocket bias enables near-native detection through Center-of-Mass drit:** Due to the pocket bias, decoy ligands are shifted further from the pocket in the protein-ligand graphs. This increases the distance between ligand and protein Center-of-Mass (COM) in these representations. **a, d)** Correlation between COM-distance and ligand RMSD under pocket bias versus individualized pocket extraction. **b, e)** Distribution shifts in COM-distance for near-native versus decoy graphs. **c, f)** ROC-AUC curves for a naive classifier using solely the COM-distance to distinguish near-native poses from decoys. The observed separability is strongly enhanced by pocket bias. Statistical annotations include p-values computed by two-sided Mann-Whitney U tests. Boxplots represent the median (center line), 25th–75th percentiles (box), and whiskers extending to 1.5 $\times$ the interquartile range (IQR). Outliers are omitted for clarity. Sample sizes ($n$) are indicated for each distribution.

### Supplementary Note 1: Size-Affinity Correlation Drives a Systematic Size Bias in the CASF2016

A size bias exists within the CASF2016 screening power benchmark: The "Top-1" ligands targeted for identification are significantly larger than the decoy ligands. This disparity artificially enhances the discrimination performance of models that positively reward molecule size.

**Curation of the Top-1 Ligands in the Benchmark** This size bias is a direct artifact of the CASF2016 screening dataset curation process. The dataset is derived from the PDBbind database, which is clustered by protein sequence similarity into 57 diverse clusters. Within each cluster, five ligands are selected to represent a wide range of binding affinities. The ligand with the highest affinity in this range is designated as the "Reference" (Top-1) ligand for that target, while the remaining four are categorized as "Other Actives." The decoys consist of 280 ligands belonging to different protein targets. Due to the inherent correlation between molecular size and binding affinity, the selection of the highest-affinity ligand leads to the Top-1 targets being larger molecules. This shift in ligand size is accompanied by a corresponding increase in the number of close protein-ligand contacts (defined as node pairs within $5\,\text{Å}$).

**The Hard-Coded Size Reward in GenScore** The GenScore architecture exploits this benchmark bias through its fundamental scoring mechanism. The model uses a siamese-style dual encoder followed by a pairwise interaction module. Instead of directly predicting a single affinity score, GenScore uses a Mixture Density Network (MDN) to predict the probability distribution of the spatial distance between every atom-residue pair. First, the ligand graph and the protein graph are passed through independent GNN encoders, resulting in updated, high-dimensional feature vectors for every ligand atom and every protein residue. Then, the two representations are combined into a pairwise interaction matrix, and a neural network predicts the probability distribution of the spatial distance between every atom-residue pair. The actual binding affinity or prediction score is derived by evaluating how well the actual 3D distances match the predicted distance distributions. For this, the true Euclidean distances are used alongside the predicted distance distributions to calculate the log-probability of those true distances occurring under the predicted distribution. These output probabilities are are summed together to produce the final predicted score for the complex.
Crucially, the scoring algorithm applies a "hard mask" to these probabilities based on the true 3D coordinates, restricting the summation to atom-residue pairs within a $5\,\text{Å}$ spatial cutoff. This is precisely where the benchmark's size bias is exploited: Because the Top-1 ligands are significantly larger and possess more $5\,\text{Å}$ contacts than the decoys, there are more probabilities to sum up. While decoy ligands are penalized with lower total scores simply due to their smaller size and fewer $5\,\text{Å}$ connections, the Top-1 ligands receive systematically higher scores due to their atom count. Consequently, the model's high screening performance in the CASF2016 benchmark is, in part, an artifact of this hard-coded size reward interacting with the size-affinity correlation of the benchmark. It is to be expected that screening performance values are lower in a realistic scenario, that is when the model is evaluated on a dataset of Top-1 ligands with similar size distribution as the accompanying decoys.

### Supplementary Figure 3: Size Bias in CASF2016 Screening Benchmark

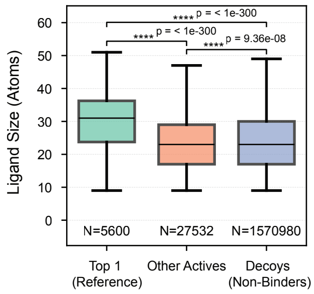


**Supplementary Figure 3:** Size bias
